# Bill color is dynamic across the breeding season but not condition-dependent in Atlantic puffins

**DOI:** 10.1101/2023.04.11.536353

**Authors:** Katja Helgeson Kochvar, Amy Catherine Wilson, Pierre-Paul Bitton

## Abstract

Sexually monomorphic species have been historically overlooked in the sexual/social selection literature, but there is growing evidence that mutual ornamentation can be driven by selective forces such as mutual sexual selection or selection for individual recognition. Examining the properties of a trait may elucidate which forces most likely play a role, especially when comparing the characteristics of quality and identity traits. Atlantic puffins *(Fratercula arctica)* are an example of a mutually ornamented monomorphic species, where both males and females display a bright orange-red bill and orange gape rosette during the breeding season and are ornamented to similar degrees. In this study, we investigate whether the properties of the colorful bill and rosette, specifically lability across the breeding season and condition-dependence, more closely align with signals of quality or identity. Our findings support prior work that the bill is sexually monochromatic from an avian visual perspective. We also determined that the bill changes in a discriminable way within individuals and is especially dynamic in the fleshy cere and rosette. However, no metric of color on any region of the bill or rosette were significantly related to current body condition. Ultimately, we argue that bill color likely functions as a quality signal, although further study is needed to determine which aspect of quality coloration signals, if not condition. These results provide a basis for experimentally testing the signal value of the colorful bill in Atlantic puffins, and more broadly, a framework for investigating the properties of mutual ornamentation in avian species.

**Lay summary:** The properties of a colorful trait inform its potential function as a signal. We investigated whether the Atlantic puffin’s colorful bill could reflect an individual’s identity or quality. Because bill color changes within the breeding season, this feature is not stable enough to reliably signal identity. Bill color is also not related to a common proxy of quality (body condition) but may communicate another aspect of quality such as foraging or parenting ability.

## Introduction

The selective forces that shape colorful displays have piqued scientific attention in recent decades. Most of this research has focused on sexually dimorphic species, in which the degree of sexual selection is presumed to be high (Andersson 1994). Yet, in many species, males and females are conspicuously ornamented to very similar degrees. In some cases, a nonadaptive genetic link between the sexes may play a role in explaining monomorphic features, as proposed by the genetic correlation hypothesis. This hypothesis posits that elaborate features are only functional in males, and are simply expressed as a genetic by-product in females (Lande 1980, 1987). However, this ignores the fact that such traits are usually costly to develop and maintain, and ultimately fails to fully explain why they are expressed in both sexes (Amundsen 2000). Indeed, there is growing evidence that selective forces also shape ornamentation in monomorphic species (Amundsen and Pärn 2006; Kraaijeveld et al. 2007; Dale et al. 2015).

Mutual sexual selection, social selection, and selection for recognition are the primary mechanisms that may explain the continued presence of elaborate displays in monochromatic species (Kraaijeveld et al. 2007). In mutual sexual selection, bidirectional mate choice or mate competition selects for an ornamented trait, whereas in social selection, competition over non-sexual resources such as territories or food resources is responsible (Kraaijeveld et al. 2007). Selection for recognition occurs when distinctive variants of a trait facilitate recognition, whether that be between species, kin, mates, or neighbors (Sherman et al. 1997; Tibbetts and Dale 2007). In mutually or socially selected traits, there is often a link between the display and some aspect of individual quality. Quality may represent a feature inherent to the signaler, such as physical condition, good genes, or age, and/or may reflect abilities acquired through experience, such as the capacity to provide parental care or secure a good territory (Dale et al. 2001). These traits are thought to be reliably maintained because low-quality individuals cannot afford the costs of elaborate trait expression, thus providing “honest” information to receivers (Zahavi 1975; Kodric-Brown and Brown 1984; Kirkpatrick and Ryan 1991; Andersson 1994). In contrast, individual recognition traits are not necessarily linked to quality, but rather are unique such that perceivers can consistently identify the informer (Dale et al. 2001; Tibbetts and Dale 2007). Individually recognizable traits can be useful in relocating mates or offspring, as well as reducing costs of territorial defense or competition in group dominance hierarchies (Tibbetts & Dale, 2007).

A trait’s features can lend insight into which selective forces play a larger role. The properties of signals of quality and signals of identity share some similarities; both types have high degrees of phenotypic variability within a population (Dale et al. 2001). However, there are also four important differences between quality and identity signals: 1) signals of quality fluctuate between and/or within years for a given individual, whereas signals of identity remain highly stable over time, 2) frequency distributions are generally unimodal for signals of quality and complex for signals of identity, 3) only signals of quality are expected to be condition-dependent and correlate with aspects of fitness, and 4) signals of quality are more heavily influenced by environmental factors, whereas signals of identity are more heavily influenced by genetic factors (Dale et al. 2001). Trade-offs between the two types of signals make it theoretically challenging for a single trait to provide both types of information, although it is still possible if different features of the trait vary independently of each other (i.e., segregation of information; Marler 1960).

Carotenoid-pigmented features have received significant attention for their role as signals of quality (Hill and McGraw 2006b). Carotenoids serve as key pigments for red, orange, and yellow features, must be acquired from the animal’s diet, and play important roles as antioxidants and immune-enhancers (Hill and McGraw 2006a). For these reasons, many hypotheses have proposed a relationship between carotenoid-pigmented features and an individual’s foraging ability (Endler 1980; Hill 1992), immune function (Lozano 1994; Moller et al. 2000), and overall health (Hill 2011). There is evidence supporting these hypotheses in some species, but there are many caveats and contradictions that make it challenging to deem carotenoid features as broadly reliable signals of quality (Svensson and Wong 2011).

Seabirds often have carotenoid-pigmented bare part coloration, standing in stark contrast to their generally achromatic plumage. Atlantic puffins (*Fratercula arctica;* hereafter: puffin) are no exception, displaying bright red-orange mandibles, an orange rosette, and orange legs and feet throughout the breeding season. The bill, which I refer to throughout this chapter as the region including the upper and lower mandibles, cere, and rosette, is very conspicuously ornamented in puffins; the difference between the colorful bill and rosette during the summer and the mostly black bill and pale rosette during the winter is stark, even to the human eye. This transition in ornamentation has been well documented in puffins (Harris and Wanless 2011), but little is known about the functional role of the colorful bill and rosette during the breeding season (see Doutrelant et al. 2013 for some discussion).

When considering the potential adaptive function of bill coloration, several aspects of puffin life history must be taken into account. Puffins are long-lived (25+ years), socially and genetically monogamous seabirds with high interannual adult survival and low divorce rates (Harris and Wanless 2011). Therefore, after initial pairing at age 4-5, most puffins stay with their mate each breeding season; only in the unlikely event of partner death or divorce do they need to select a new mate (Harris and Wanless 2011). Additionally, puffins exhibit obligate bi-parental care of a single chick, with equal levels of care provided by males and females (although roles differ slightly; Creelman and Storey 1991). It is thus unsurprising that puffins are sexually monochromatic (accounting for avian vision; Doutrelant et al. 2013), which classically would indicate that puffins experience low levels of intersexual selection.

However, the potential for mutual sexual selection, social selection, or selection for recognition in shaping the colorful bill should not be ruled out. Mutual sexual selection may operate on dynamic assessment traits used to make decisions concerning parental investment (Kraaijeveld et al. 2007), especially in species like puffins where there is a strong trade-off between current reproductive effort and future survival/breeding attempts (Trivers 1972; Johnsen et al. 1994; Erikstad et al. 2009). Social selection may operate when individuals engage in fierce, at times physically aggressive competition for burrows at the beginning of the breeding season (Harris and Wanless 2011). Additionally, puffins have high interannual mate and site fidelity, so it is highly plausible that social networks remain consistent from year to year. Individual recognition could be especially useful in reuniting with a long-term mate or limiting costly territorial defense between familiar neighbors.

While the function of bill color and the selective forces that shape it remain unknown, knowledge of its properties can be used to generate more informed hypotheses. In Atlantic puffins, bill and rosette color may 1) communicate some aspect of individual quality (between mates or to conspecifics), and/or 2) provide information on individual or kin identity. A central aim of this study is to determine whether puffin bill coloration more closely corresponds to the properties of a quality signal or an identity signal, or alternatively, whether some aspects of the bill resemble quality signals while other aspects resemble identity signals. Specifically, we will focus on the lability and condition-dependence of bill coloration. The degree to which Atlantic puffin bill colouration changes is unknown, but could theoretically fluctuate within a few days, as in other species with carotenoid-pigmented bills (Ardia et al., 2010; Rosenthal et al., 2012). The condition-dependence of bill colouration has been previously investigated in puffins, but with conflicting results (support from Doutrelant et al., 2013; lack of support from Kelly, 2015).

This paper lays the foundation for exploring the adaptive value of color in the puffin’s bill, cere, and rosette. First, we seek to replicate Doutrelant et al.’s (2013) key finding that puffins are sexually monochromatic on the bill. Doutrelant et al. (2013) assessed the dichromatism of two regions on the bill and the rosette with a non-parametric multivariate model to compare the colour space occupied by the two sexes, as well as a generalized linear model with the four photoreceptor responses of each region as predictors of sex. Neither analysis yielded significant results, providing evidence that these regions are monochromatic from an avian perspective (Doutrelant et al., 2013). Our analysis will build on this work by 1) employing both statistical and perceptual approaches to evaluate dichromatism, 2) incorporating a robust sample size of hundreds of individuals (compared to 36 in Doutrelant et al., 2013), and 3) sampling the cere in addition to the bill and rosette. We then characterize key properties of puffin bill coloration (lability and condition-dependence) to evaluate the bill’s potential role as a quality signal and/or identity signal, and whether mutual sexual selection, social selection, or selection for individual recognition are more likely to be acting.

## Methods

### Study site

This study was conducted on Gull Island in the Witless Bay Ecological Reserve of Newfoundland and Labrador, Canada (47.26, -52.77). The Atlantic puffin colony on Gull Island is one of the largest in the Northwestern Atlantic, with ∼120,000 breeding pairs according to a 2012 population survey (Wilhem 2017).

### Field methods

To assess whether Atlantic puffin bill coloration changes within a breeding season, we employed a cross-sectional (across individuals) and longitudinal (within-individual) approach. For the cross-sectional analysis, as well as the analysis of sexual dichromatism, we used morphometric and color data from 229 individuals, sampled during late June to mid-August of 2019, 2020, and 2021. Some individuals were sampled multiple times within or between years, which we account for in our statistical analyses (see Statistical analyses). We focused on the brood rearing part of the breeding season because puffins are particularly sensitive to human disturbance during the incubation phase, and are much less likely to abandon after the chicks have hatched (Rodway et al. 1996). The mean hatch date in this population tends to be in the last week of June or first week of July, so we began sampling after many adults were no longer incubating. In addition, when extracting puffins from their burrows, we assessed whether there was an egg or chick present and did not further handle the adult if an egg was identified. To assess color changes longitudinally, we opportunistically sampled 41 individuals twice within a year during late June to mid-August of the years 2019, 2020, and 2021. The two sampling dates took place 1-28 days apart, with a mean of 12.02 ± 5.22 days.

Individuals were captured by hand in their burrows between the hours of 22:00 and 3:00. At first capture, individuals were given a Canadian Wildlife Service stainless steel band for subsequent identification, mass was measured with a 600 g Pesola spring scale to the nearest 5 g, flattened wing chord was measured with a ruler to the nearest 1 mm, and a blood sample was taken from the brachial vein on #2 filter paper for sex determination. Individuals were then taken into a blind, where auto-exposure bracketed ultraviolet (UV, 320-380 nm) and visual spectrum (400-680 nm) RAW 20 megapixel images were taken of the left side of the beak and rosette with a full spectrum converted Samsung NX1000 (following instructions from Troscianko, 2018) using two 2-inch Baader lens filters (Fig. 1). UV and visual spectrum images were obtained because unlike humans with three photoreceptors, puffins have a fourth photoreceptor sensitive to UV wavelengths (Endler and Miekle 2005), and our goal was to measure bill colors as they would be perceived by a puffin. The photos were illuminated with a full-spectrum Metal Halide 150W ballast that passed through a light diffuser, and all photos included optic grade white (99% reflectance) and dark (10% reflectance) standards, a ruler, and a small white board to record the date and band number of the individual puffin. The bill was cleaned of debris with a toothbrush prior to photo capture and was held in place with a wooden bill stabilizer. Handling time for these procedures was typically no more than 15 minutes, and in no case did a puffin exhibit typical signs of distress (i.e., panting). This process was repeated for the second capture, except that redundancies were avoided by excluding the need to band the bird, measure the wing chord (it remains relatively stable throughout the breeding season), and collect blood (only one sample needed for sexing).

**Figure 1.**
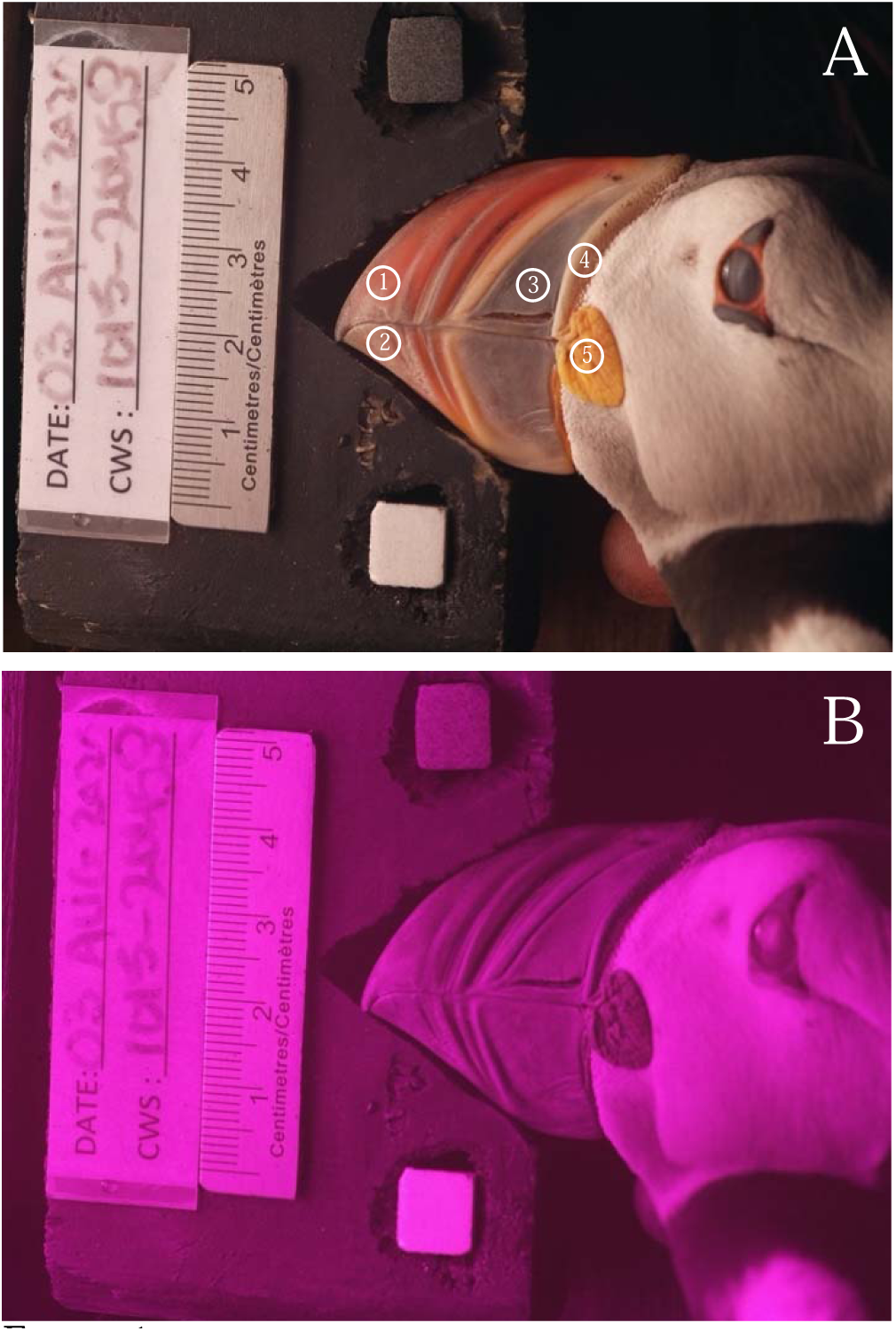
Example photos of an adult puffin from 2020 following the steps outlined in the methods. Each puffin was photographed in the A) visual spectrum and B) ultraviolet spectrum. The visual spectrum photo (A) also displays the locations of regions of interest (ROIs). ROIs were selected for each individual puffin on the 1) upper mandible, 2) lower mandible, 3) base of the mandible, 4) cere, and 5) rosette. Selection areas are enlarged on the illustration to show detail.

### Molecular methods

Sex of each individual was determined molecularly from blood samples collected in the field. Blood was stored at room temperature until molecular procedures could begin. At this time, a ∼1 cm^2^ section of paper saturated with blood was extracted with sterilized scissors and placed in a 1.5 ml collection tube. DNA was extracted using a DNeasy® Blood & Tissue Kit (Qiagen Inc., Toronto, ON, CA) following protocols outlined in the DNeasy® Blood & Tissue Handbook (2020) and stored at -20 °C.

Polymerase chain reaction (PCR) was run with extracted DNA to amplify the chromo-helicase DNA 1 (CHD1) gene on the avian W and Z chromosomes. Following standard procedures for sexing seabirds (Fridolfsson and Ellegren 1999), 12.5 µl Thermo Scientific_™_ PCR Master Mix, 2 µl of both primers 2550F and 2718R, 6.5 µl of nuclease-free water, and 2 µl of extracted DNA were added to each PCR tube. All batches were run with a no template control (NTC) tube, which contained an additional 2 µl of nuclease-free water instead of extracted DNA. The PCR was performed with an Eppendorf Mastercycler® ep gradient S on a program of 95 °C for 5 minutes, 35 cycles of denaturing, annealing, and extension at 94 °C for 30 seconds, 50 °C for 30 seconds, 72 °C for 60 seconds, extension at 72 °C for seven minutes and a cooling period of 4 °C for 10 minutes. PCR samples were stored at -20 °C unless proceeding directly to gel electrophoresis.

PCR samples were run on a RedSafe_™_ agarose gel with 100 base pair reference ladders on a Thermo Scientific_™_ EC 300 XL for 50 minutes at 130 amps. The gels were then imaged using Image Lab software and stored digitally. All procedures were carried out at Memorial University of Newfoundland following standard lab safety protocols.

### Calculation of body condition index

Body condition was determined from the residuals of a best fit linear regression with mass as the response variable. Typically, body condition is either given as the residuals of a linear regression of mass on wing chord length or simply mass. However, mass in puffins is known to change across the breeding season, vary due to fluctuations in prey availability and environmental conditions, and differ between the sexes, so these variables (Julian date of capture, year of capture, and sex) were included in the regression model. To account for potential sex-biased differences in mass change within and between years, we also included the two-way interactions between sex and Julian date, sex and year, as well as the three-way interaction between sex, Julian date, and year in the full model. Prior to fitting the model, mass and wing length were centered by subtracting the mean value to reduce instances of structural multicollinearity between main-effect and interaction terms (Cohen et al. 2014). The full model was reduced by stepwise removal of non-significant terms based on results from ANOVA tables. The final linear regression model met the assumptions of homogeneity of variance, normal distribution of the residuals, and low multicollinearity among predictors.

### Assessment of color

Multispectral images of the left side of the beak were generated using micaToolbox 2.2v in ImageJ for each individual at a given capture date (van den Berg et al. 2020). One visual spectrum and one UV RAW photo were chosen from the two sets of bracketed photos for alignment using the Photoscreening tool, permitting selection of the most illuminated sample without evidence of RGB camera pixel oversaturation. Visual spectrum and UV images of the puffin bill were aligned and merged with either the ‘Affine align’ tool by selecting four consecutive points on the bill of each image (uppermost intersection of upper mandible and cere, bill tip where mandibles meet, bottommost intersection of lower mandible and cere, intersection of cere and rosette), or the ‘Manual align’ tool by manually positioning the photos such that the bills exactly match up. The quality of the alignment was evaluated by creating a false color image with the ‘Make Presentation Image’ tool, using yellow to represent the visual R normalized channel and blue to represent the uvB normalized channel. False color images without evidence of poor alignment on the bill were considered aligned and could be used in subsequent analysis. From these multispectral images, cone catch images were generated using the spectral sensitivity of a violet-sensitive (VS) avian visual system (λ = 410, 450, 505, 565 nm; Ödeen et al. 2010). The spectral sensitivity of puffin vision is currently unknown, but other closely related species in the family *Alcidae* (i.e., common murres *Uria aalge* and razorbills *Alca torda*, Ödeen et al. 2010) are VS based on sequencing of the SWS1 opsin gene. Additional parameters in the cone catch model included double cones representing perception of brightness and standard illuminant D65 (CIE) as a typical spectrum under ambient light conditions. Quantum catch values for five regions of interest (ROIs) on the bill were extracted from the cone catch images: two on the tip of the upper and lower mandible, where puffins generally have the most saturated carotenoid-pigmented coloration, one on the base of the mandible, which is black and presumably melanin-pigmented, one on the semi-fleshy cere, which should also respond rather dynamically to changes in carotenoid levels, and one on the fleshy rosette, which should respond most dynamically to fluctuations in carotenoid availability (Fig. 1). Each ROI was the same shape and size (30x30 pixel circle) and positioned to obtain the clearest measure of color possible (i.e., avoiding obvious scratches or pieces of dirt). Average quantum catch values from the ROIs were modelled in tetrachromatic color space using the ‘colspace’ function, with space set to tetrochromatic (“tcs”) and qcatch set to quantum catch (“Qi”; Maia et al., 2013). A tetrahedral color space defines a 3-dimensional space of perceivable colors, where the central point is the achromatic center, representing equal stimulation of the cones, and each vertex is the absolute stimulation of one of the four cones. Colors from ROIs plotted in this space have Cartesian coordinates (x,y,z) and spherical coordinates (0, <, r), the latter of which was used in both the cross-sectional and longitudinal analyses.

From the spherical coordinates, a color vector can be defined between the achromatic center and the position of the measured color in color space. The color vector can be subsequently used to identify two measures of hue and one measure of chroma for each ROI. Hue is the direction of the color vector (drawn from the achromatic center to the locus) and is defined by the angular position of the locus in tetrahedral color space, both in terms of the azimuth (horizontal axis, 0) and elevation (vertical axis, ¢; Stoddard & Prum, 2008). Hue 0 (hereafter “hue VIS”) represents the contribution of the visible spectrum (red-blue) to perceived color. Hue VIS ranges from -π to +π, such that perceived reds and purples have negative values, perceived yellows and oranges are close to zero, and perceived greens and blues have positive values (Dakin and Montgomerie 2013). Across the ROIs in our study sample, hue VIS ranged from red-orange (-0.354) to orange-yellow (0.507). Hue ¢, hereafter “hue UV”, represents the contribution of violet and UV to perceived color. Hue UV ranges from -π/2 to +π/2, with more UV rich colors having more positive values (Dakin and Montgomerie 2013). UV reflection is low across the measurements in our study sample, with the maximum value on the upper mandible (-0.375), the minimum value on the cere (-1.170), and a -0.704 average across the regions. Chroma is the saturation of a color and is defined as the magnitude of the color vector (*r*) from the achromatic center (Stoddard and Prum 2008). Because the color space is a tetrahedron, different hues have different potential maximum chroma (*r_max_*), so achieved chroma (*r/ r_max_*) was calculated for each ROI as a more informative estimate (Stoddard and Prum 2008). These measures were determined for all four chromatic regions (upper and lower mandible, cere, and rosette).

Luminance was calculated from the relative stimulation of the double cone as a measure of brightness. Brightness is the sole achromatic measure and represents how much light is reflected from the surface. Brightness was measured for all five regions, including the achromatic mandible base. All colorimetric variables were calculated with the R package *pavo* (Maia et al. 2013, 2019).

Because these colorimetric variables do not give us information on the perceptual aspect of color (i.e., discriminability), a modeling approach was employed to statistically quantify the degree of sexual dichromatism in the population, as well as to evaluate longitudinal differences in color within a breeding season. Rather than working with Cartesian or spherical coordinates, we directly transformed the quantum catch values using the ‘coldist’ function in the R package *pavo* (Maia et al. 2013, 2019). The noise-weighted Euclidean distance between the two sets of quantum catch values for each individual was calculated using the receptor noise-limited (RNL) model, which estimates color and brightness discrimination thresholds based on receptor noise (Vorobyev and Osorio 1998). Chromatic differences (dS) were calculated from the quantum catch values of the four single cone channels, whereas achromatic differences (dL) were calculated from the quantum catch value of the double cone channel. Receptor noise was estimated from relative densities of photoreceptor types, which we estimated as 1:1:3:3.55 (UV-wavelength sensitive, short-wavelength sensitive, medium-wavelength sensitive, and long-wavelength sensitive photoreceptors, respectively) for Atlantic puffins based on the proportion of oil droplet types in the retina (Émond 2016). The Weber fraction, which is used to determine the signal-to-noise ratio, was set at 0.10 for the chromatic channel and 0.15 for the achromatic channel based on average results from behavioral tests of other avian species (Olsson et al. 2018). Noise-weighted Euclidean distances generated from this model correspond directly to perceptual differences (termed just noticeable differences, JNDs; Vorobyev et al. 1998; Vorobyev and Osorio 1998). The general rule is that a chromatic or achromatic difference is likely perceptible to conspecifics if the JND > 1, whereas two colors should be indistinguishable if the JND < 1. However, this is often only the case under optimal viewing conditions (i.e., controlled laboratory studies), so JND values of two and three may be more relevant to animal communication in a natural setting, where illumination changes, environmental distractions, and temporal constraints may detract from the ability to reliably discriminate colors (Fleishman et al. 2016).

### Statistical analyses

#### Sexual dichromatism

To evaluate whether puffins are sexually dichromatic on the bill, cere, and rosette, we calculated whether each bill region was statistically separate in color space between males and females, as well as perceptibly discriminable. Following the proposed methodology outlined by Maia and White (2018), statistical separation was evaluated using noise-corrected color and brightness distances (generated with the ‘coldist’ function) to run a distance-based PERMANOVA (hereafter: distance PERMANOVA) with the ‘adonis’ function in the R package *vegan* (Oksanen et al. 2007). Distance PERMANOVA is a nonparametric test that simulates a null distribution by randomizing distances between observations, resulting in the generation of a pseudo-F statistic (Anderson 2017; Maia and White 2018). We ran this on each ROI with 999 permutations to determine whether the region was significantly different in color and/or brightness space between the sexes (α = 0.05) and estimate the effect size of the analysis (*R*^2^). *P* values were adjusted across all distance PERMANOVA tests with the false discovery rate to correct for type 1 error (Benjamini and Hochberg 1995).

Perceptible discrimination was evaluated using color and brightness distances generated from a bootstrapping procedure with the ‘bootcoldist’ function in *pavo* (see section on Assessment of color for ‘coldist’ settings). This function samples with replacement 1000 times (“boot.n” = 1000) to generate a distribution of mean distances between the two groups, in this case males and females. From this, 95% confidence intervals around the geometric mean distance could be estimated. Distances with 95% confidence intervals above 1 JND were considered theoretically discriminable, whereas those with intervals above 3 JND were considered reliably discriminable across sensory environments. Both thresholds are useful in assessing the degree to which males and females are dichromatic from a puffin’s perspective. For both the statistical and perceptible dichromatism analyses, 162 of 229 samples were retained (77 male, 85 female), as they represented unique observations of genetically sexed individuals.

#### Trends in color change over the breeding season

The cross-sectional dataset was used to assess broad temporal trends in coloration across the late breeding season. Linear models were generated for each colorimetric variable, where hue, chroma, or brightness was the response variable and Julian date was the sole predictor variable. Diagnostic plots were generated to ensure the residuals met the assumptions of normality, linearity, and homoscedasticity. For individuals that were sampled more than once within and/or between years, only one of the sampling dates was retained in the dataset. This was randomly chosen using an R sampling procedure, such that some regions from the same individual retained values from different sampling dates. There were 53 instances where individuals were sampled multiple times within or between years, so 176 of 229 observations were included in each model.

#### Discriminability

The longitudinal dataset was used to assess whether individuals perceptibly change color within the breeding season. To assess the scope of discriminable changes, the percentage of regions that changed color or brightness in a discriminable vs. non-discriminable way within individuals was calculated for each ROI separately at JND values of one, two and three. Achromatic differences were calculated for all five ROIs, whereas chromatic differences were calculated for four of the five ROIs, excluding the black, achromatic base of the mandible. These percentages give a general assessment of whether puffin bill coloration perceptibly fluctuates during the breeding season.

We also investigated the effect of time between sampling on the discriminability of color or brightness using unpaired t-tests, with discriminability at a JND value of one as the dependent variable (categorical, 2 levels) and the number of days between sampling as the independent variable (continuous). To determine if the data met the assumptions of normality and equal variance, a Shapiro-Wilk’s test of normality and a Levene’s test of homogeneity of variance was run on the original discriminability group distributions. If these assumptions were not met (α = 0.05), a Mann-Whitney U test was performed. Outliers were identified with boxplots, and for parametric t-tests only, one outlier was removed when analyzing each region separately because it fell well outside of the whiskers of the boxplot (i.e., more than 1.5x the inter-quartile range). The magnitude of the difference in days between sampling for discriminable vs. indiscriminable colors was calculated based on the group medians and reported alongside interquartile ranges (Q1, Q3). Effect sizes for t-tests or Mann-Whitney U tests were also calculated and interpreted using Cohen’s cut-offs (small: 0.1 –< 0.3, moderate: 0.3 –< 0.5, large: >= 0.5). Finally, a linear regression was used to predictively assess the temporal threshold for color and brightness discriminability of each region, with change in Euclidean distance (measured in JNDs) as the response variable (y) and time difference (in days) as the predictor variable (x). The regression equation was reverse solved to determine the threshold for discriminability (i.e., when the regression equation equals one JND).

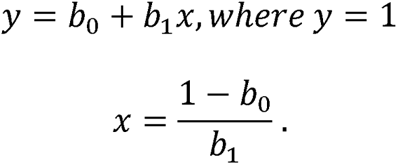

This equation was solved for all individuals, as well as males and females separately, to evaluate sex-biased discriminability differences. The mean and standard error of the threshold was calculated by sampling the variables *b*_0_ and *b*_1_ from their respective ranges (estimate ± standard error) with replacement 1000 times. Only the thresholds of significant linear relationships after correction for type I error with a false discovery rate are reported (Benjamini and Hochberg 1995).

#### Color and time interval between sampling

The longitudinal data was also used to determine if changes in color over the breeding season are due to individual changes over time. Specifically, the relationship between change in coloration/brightness and the number of days elapsed between sampling dates was assessed with linear models, controlling for date of first capture. The interaction between time difference and date of first capture was also included in the initial model and retained if the interaction was significant. This was analyzed for each ROI and each colorimetric variable separately. Change in coloration and brightness was assessed as the raw difference between these values on successive sampling dates (C_2_ – C_1_, where C_2_ is the value at second capture and C_1_ is the value at first capture). Positive values indicated that the metric increased, whereas negative values indicated that the metric decreased. Diagnostic plots were produced for each model to determine if the residuals met the assumptions of normality, linearity, and homoscedasticity. Cook’s distance was used to identify influential outliers by removing values that had a Cook’s distance value four times greater than the mean. All models met the assumptions of linear regression after removal of outliers.

#### Color and condition

The relationship between color and condition was evaluated with the cross-sectional dataset using linear models, where hue, chroma, or brightness were the response variables and body condition was the predictor variable. Because Julian date of capture, year, and sex were included in the regression from which body condition was calculated, these variables were not included as separate predictor variables in the models. Diagnostic plots were produced for each model to determine if the residuals met the assumptions of normality and homoscedasticity. Cook’s distance was used to identify influential outliers by removing values that had a Cook’s distance value four times greater than the mean. For individuals that were sampled multiple times within or across years, only one observation was randomly chosen to be retained in the dataset, yielding a final dataset of 162 individuals.

## Results

### Sexual dichromatism

After applying the false discovery rate correction, the distance PERMANOVA showed that male and female coloration was not statistically different in color or brightness space on any region (*P >* 0.05, Table S1, Table S2).

All the chromatic bootstrapped estimates had 95% confidence intervals either completely below 1 JND (upper mandible, lower mandible, base of mandible, cere), or overlapping with 1 JND (rosette; Fig. S1). For the achromatic bootstrapped estimates, the base of the mandible was the only region that overlapped with 1 JND (Fig. S1). Neither the chromatic nor achromatic bootstrapped estimates approached 3 JND. Based on these estimates, none of the five bill regions are perceptibly different between males and females, even under ideal conditions. Importantly, our results are not influenced by differences in sampling between the sexes, as there was no statistical difference between the capture dates of males vs. females (t = 0.94, *P* = 0.347).

### Color trends over the breeding season

The cross-sectional dataset was used to evaluate broad trends in color change across the breeding season. We found that several colorimetric variables in different regions of the bill, cere, and rosette changed within the breeding season (Table 1):

1. Upper mandible measures of hue and brightness significantly changed over the breeding season. Brightness decreased and the hue became more orange and less UV-reflective (Table 1; Fig. 2A, B, D)
2. Lower mandible saturation significantly increased over time, whereas brightness significantly decreased over time (Table 1; Fig. 2C, D). Neither measure of hue significantly changed over the breeding season (Table 1).
3. Base of mandible did not significantly change across the breeding season (Table 1). Measures of hue and chroma are not relevant to black features like the base of the mandible and thus were not tested.
4. Cere saturation and the UV component of hue significantly increased over the breeding season (Table 1; Fig. 2B, C). Hue VIS and brightness did not change in a significant way (Table 1).
5. Rosette measures of hue significantly changed over the breeding season, such that the rosette became more yellow and less UV reflective later in the season (Table 1; Fig. 2A, B). Brightness significantly increased, whereas achieved saturation remained unchanged (Table 1; Fig. 2D)

**Table 1.** Change in colorimetric variables across the breeding season Response: colorimetric

| ROI | Response: colorimetric variable | Estimate | Std. Error | t | P |
| --- | --- | --- | --- | --- | --- |
| Upper mandible | **Hue vis** | 0.0007 | 0.0003 | 2.222 | 0.047* |
|  | **Hue UV** | -0.0015 | 0.0006 | -2.474 | 0.030* |
|  | Achieved saturation | 0.0013 | 0.0006 | 2.000 | 0.062 |
|  | **Brightness** | -0.0013 | 0.0006 | -2.280 | 0.045* |
| Lower mandible | Hue vis | 0.0006 | 0.0004 | 1.728 | 0.104 |
|  | Hue UV | 0.0000 | 0.0006 | 0.033 | 0.974 |
|  | **Achieved saturation** | 0.0023 | 0.0006 | 3.928 | <0.001*** |
|  | **Brightness** | -0.0024 | 0.0006 | -3.908 | <0.001*** |
| Mandible base | Brightness | 0.0002 | 0.0004 | 0.543 | 0.624 |
| Cere | Hue vis | -0.0008 | 0.0004 | -2.065 | 0.062 |
|  | **Hue UV** | 0.0019 | 0.0005 | 3.891 | <0.001*** |
|  | **Achieved saturation** | 0.0024 | 0.0007 | 3.356 | 0.002** |
|  | Brightness | -0.0015 | 0.0008 | -2.015 | 0.062 |
| Rosette | **Hue vis** | 0.0024 | 0.0005 | 5.068 | <0.001*** |
|  | **Hue UV** | -0.0018 | 0.0003 | -6.174 | <0.001*** |
|  | Achieved saturation | -0.0004 | 0.0006 | -0.661 | 0.577 |
|  | **Brightness** | 0.0058 | 0.0013 | 4.338 | <0.001*** |

*P* values corrected with false discovery rate method. Significant variables bolded at *P* = 0.05\*, 0.005\*\*, 0.0005\*\*\*

**Figure 2.**
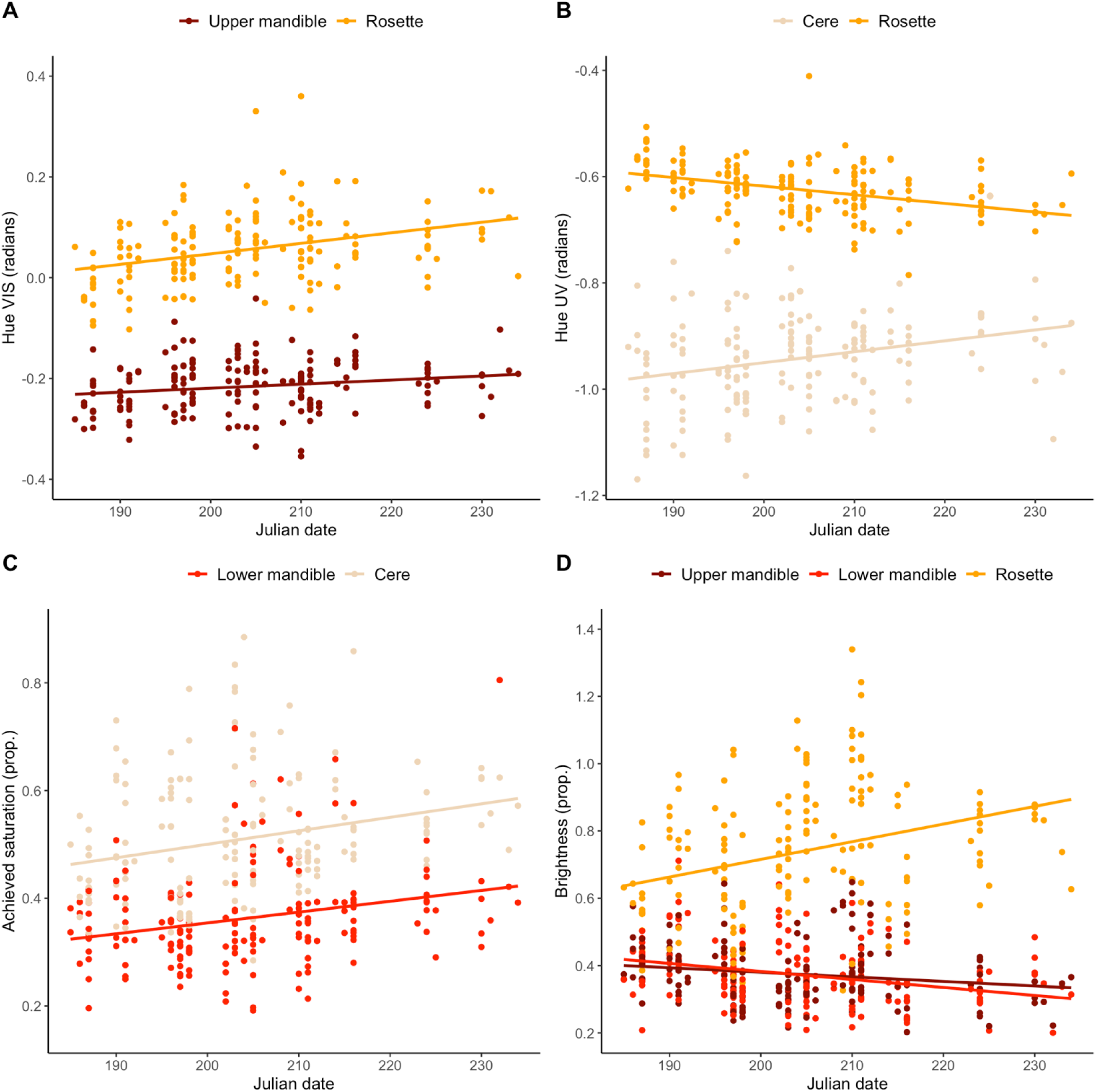
Across the late breeding season, significant changes were observed in A) hue VIS of the upper mandible and rosette, B) hue UV of the cere and rosette, C) achieved saturation of the lower mandible and cere, and D) brightness of the upper and lower mandible and rosette. Significance was obtained from linear regressions, and only significant trends after correction for the false discovery rate are displayed.

The effect of sex on these relationships was evaluated with linear models, where the main effects of capture date and sex, as well as the interaction between date and sex, were included as predictor variables. None of these relationships varied between males and females, as the interaction term was nonsignificant for all of the colorimetric variables after controlling for multiple testing with the false discovery rate (*P* > 0.05).

### Perceptible changes in coloration

The cross-sectional data was also used to evaluate whether color changes in a perceptible way within individuals. Overall, slightly more than half of the changes in coloration on the bill, cere, and rosette were discriminable based on a model of puffin vision (56.7%, Table S3; Fig. S2; JND = 1). This was driven mostly by changes in the rosette, where 80% of color changes were considered discriminable. This contrasts sharply with the lower mandible, where only 34% of changes were discriminable. As expected, fewer color changes were discriminable at higher values of JNDs; at JND = 2, less than a quarter of changes were discriminable (22.0%), and at JND = 3, less than 10% of all color changes were discriminable (8.5%, Table S3).

The number of days between sampling dates only weakly predicted whether the observed colour change was discriminable. The assumption of homogeneity of variance was met for all regions, but several did not meet the assumption of normality, in which case Mann-Whitney U tests were performed. Across all regions, the changes that were discriminable at JND = 1 had significantly more days between capture dates than changes that were considered non-discriminable (W = 2707.5, *P =* 0.046). However, the median number of days was the same for both discriminability groups (discriminable: *Mdn* = 13.0, IQR = 7.5-14.0; indiscriminable: *Mdn* = 13.0, IQR = 8.0-15.0), and the effect size was low (r = 0.156). The rosette exhibited a much stronger contrast (W = 59.5, *P =* 0.017, r = 0.376), such that discriminable changes had five more days between sampling dates (discriminable: *Mdn* = 14.0, IQR = 12.0-15.0; indiscriminable: *Mdn* = 9.0, IQR = 4.25-13.0). For all other bill regions (i.e., upper and lower bill mandible, and cere), this relationship was nonsignificant.

The threshold of discriminability could only be calculated for the rosette, representing the only significant relationship between a region’s color difference and time difference (Fig. 3). The regression neared significance for both males and females together (*t*_1,39_*=* 2.73, *P =* 0.057) and was significant for females only (*t*_1,18_ *=* 3.96*, P =* 0.011). The threshold was calculated at 4.49 ± 0.98 days for males and females (at JND = 1) and 7.79 ± 0.07 days (at JND = 1) for females only. None of the linear relationships between color difference and time difference were significant for males and females separately.

**Figure 3.**
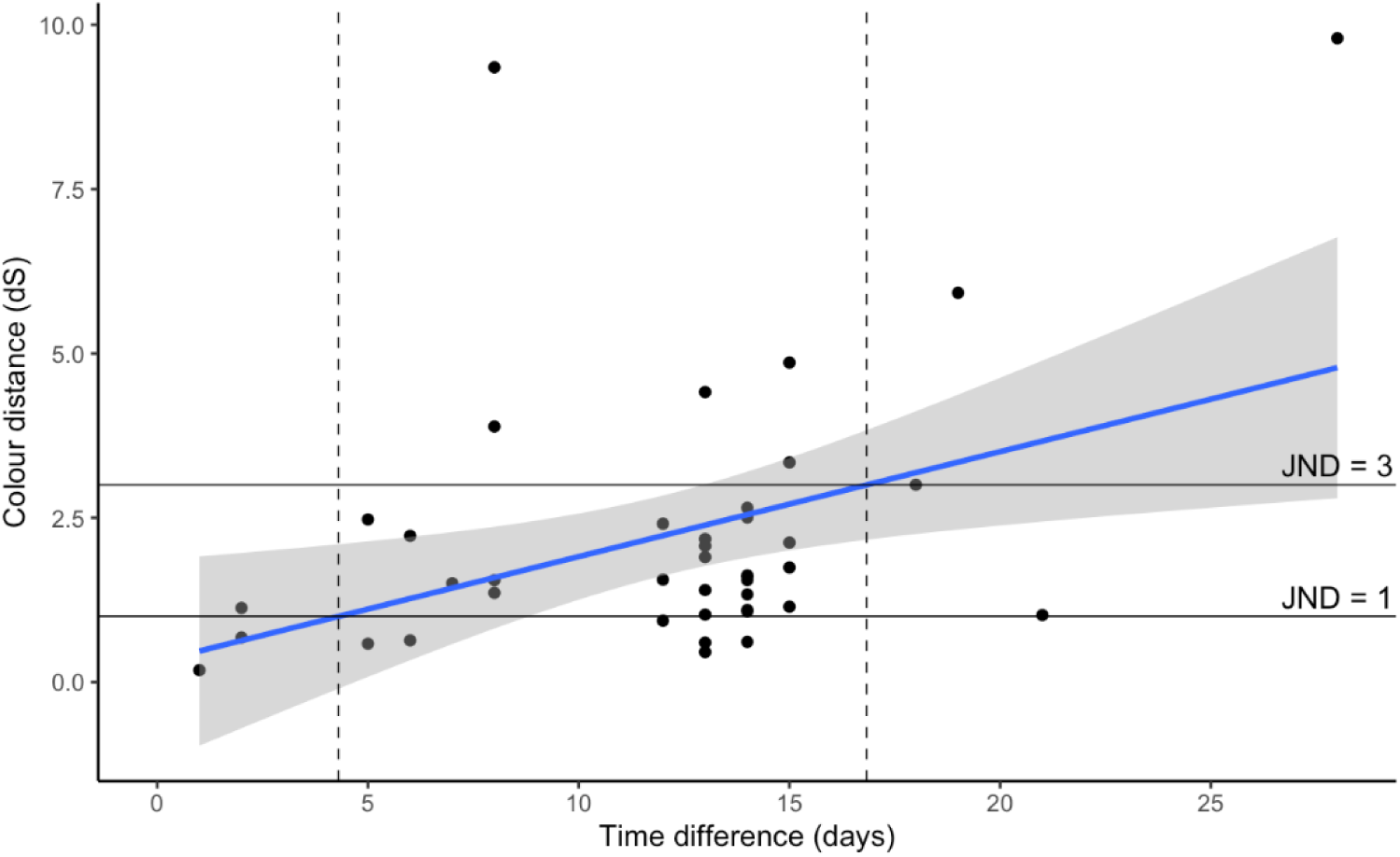
The relationship between rosette color distance and time between sampling was significant, permitting calculation of thresholds of discriminability at JND = 1 (4.30 days) and JND = 3 (16.83 days).

### Perceptible changes in brightness

Over 70% of all changes in brightness were discriminable (71.7%, Table S4, Fig. S3), and the majority of changes in all five regions were discriminable (upper mandible: 78.0%, lower mandible: 58.5%, base of mandible: 73.2%, cere: 61.0%). As with changes in coloration, the rosette had the highest proportion of discriminable brightness changes, with 88% of brightness changes considered discriminable (87.8%, Table S4).

Mann-Whitney U tests were performed to assess the relationship between discriminability and time between sampling, since the assumption of normality was violated in all cases. There was a highly significant difference in the number of days between sampling dates for discriminable (*Mdn* = 13.0, IQR = 12.0 - 15.0) vs. non-discriminable changes (*Mdn* = 12.0, IQR = 7.0-14.0), although the difference was small and the effect size was low (W = 1971.5, *P* = 0.004, r = 0.223). The rosette was the only individual region with a significant difference and a moderate effect size, where discriminable changes had about six and a half days more between sampling dates (discriminable: *Mdn =* 7.0, IQR = 5.0-8.0; indiscriminable: *Mdn* = 13.5, IQR = 12.0-14.25; W _=_ 28.5, *P=* 0.014, r = 0.387).

The threshold of discriminability (JND = 1) was not calculated for any of the five regions of the bill because no significant relationships were detected between brightness difference and time difference. When examined separately for each sex, two relationships were initially significant for males only (base of the mandible, *t*_1,18_ *=* 2.54, *P =* 0.021; rosette, *t*_1,18_ *=* 2.76, *P =* 0.013), but were nonsignificant after correcting for multiple testing with the false discovery rate (base of the mandible, *t*_1,18_ *=* 2.54, *P =* 0.154; rosette, *t*_1,18_ *=* 2.76, *P =* 0.154).

### Longitudinal changes in colorimetric variables

After applying the false discovery rate error correction, none of the colorimetric variables were significantly influenced by the number of days between sampling (Table S5). Sampling interval was initially significant for cere hue VIS, such that the cere became more orange with increasing sampling interval. Interestingly, the interaction between hue VIS and date of capture was also significant, such that the opposite trend (i.e., significant decrease with sampling interval) was observed for individuals captured later in the season (Fig. 4). These results should be interpreted with caution, however, as all terms in the model were nonsignificant after application of the false discovery rate error correction. Achieved saturation of the cere and brightness of the rosette were the only colorimetric variables that approached significance with the false discovery rate correction (α < 0.1). As the time between capture dates increased, the cere’s achieved saturation and the rosette’s brightness decreased (Fig. 4).

**Figure 4.**
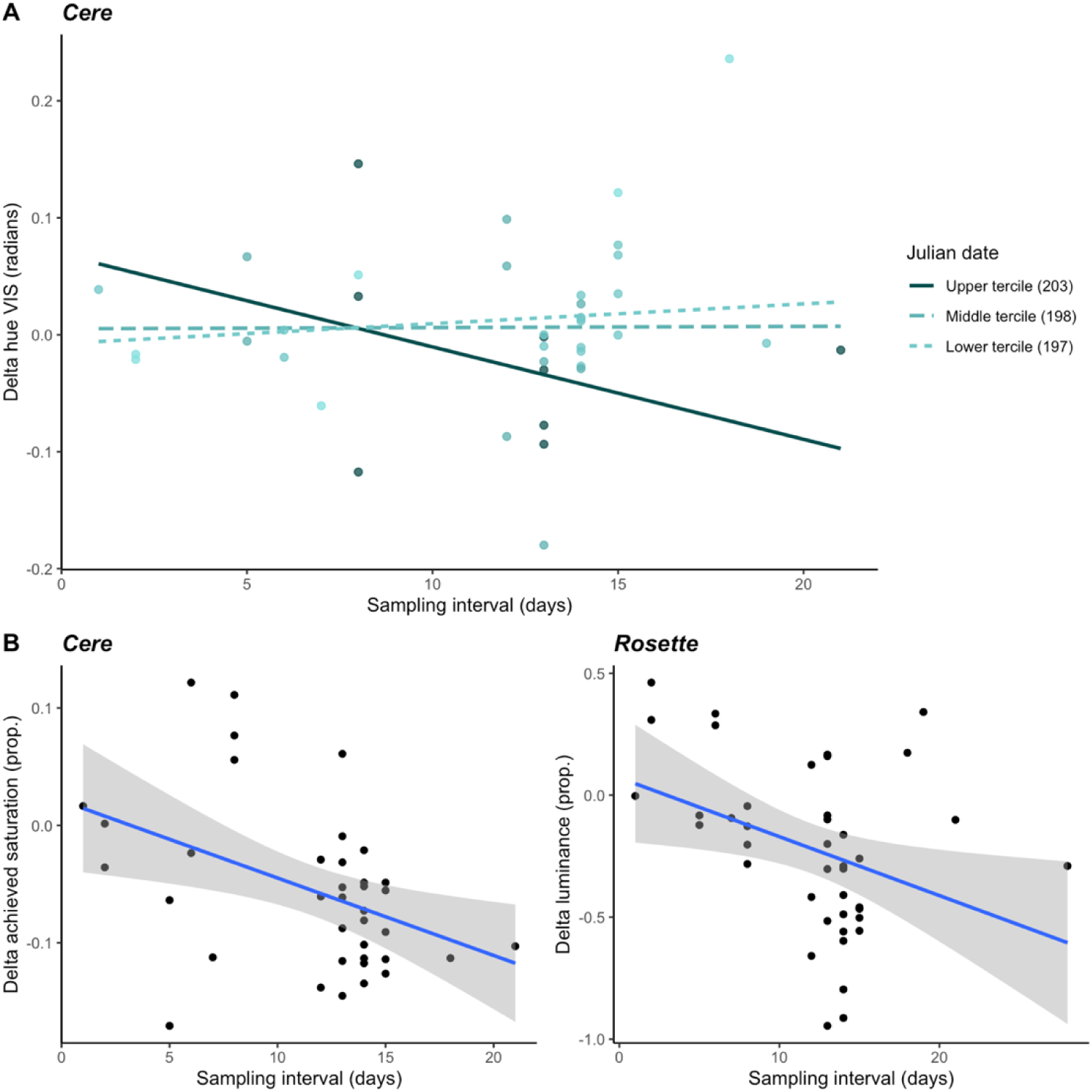
Several trends were observed regarding the relationship between change in coloration and the time elapsed between sampling dates. A) The two-way interaction between delta hue VIS and sampling interval was significant for the cere prior to application of the false discovery rate. Each line represents predictions for the mean Julian date of three equally sized terciles. For individuals captured later in the season (upper tercile), cere hue VIS negatively changes (i.e., becomes yellower) with increases in sampling interval. In contrast, cere hue VIS slightly positively changes (i.e., becomes more orange) with increases in sampling interval for individuals captured earlier in the season (lower tercile). B) Larger decreases in the cere’s achieved saturation and in the rosette’s brightness were observed with increased sampling interval.

### Color and condition

Body condition was calculated based on the residuals of a linear regression model with all four main-effect variables: wing length (scaled), Julian date of capture, year of capture, and sex. Mass was positively related to wing length (2.02 ± 0.43, F_1,162_ = 49.83, *P* < 0.001), but negatively related to Julian date (-0.73 ± 0.21, F_1,162_ = 11.39, *P* < 0.001). Mass was also higher in males compared to females (34.21 ± 3.63, F_1,162_ = 88.59, *P* < 0.001). Additionally, a post-hoc assessment of estimated marginal means found that mass was significantly lower in 2022 compared to 2019 (14.59 ± 5.14, t_162_ = 2.84, *P* = 0.026) or 2020 (16.22 ± 5.64, t_162_ = 2.88, *P* = 0.024). All the included variables were highly significant (wing length: *F*_1,162_ *=* 49.8, *P* < 0.001; Julian date of capture: *F*_1.162_ *=* 11.4, *P* < 0.001; year of capture: *F*_3,162_ *=* 5.5, *P* = 0.001; sex: *F*_1,162_ *=* 88.6, *P* < 0.001) and no evidence of multicollinearity (VIF < 1.3) was detected in the final model.

Hue VIS on the lower mandible was significantly positively related to condition before correcting for multiple testing (t_1,160_ = 2.344, *P* = 0.020). However, after application of the false discovery rate correction, no significant relationships were detected between any of the metrics of color and condition (Table S6). Assumption checks identified multiple outliers (three for upper mandible achieved saturation, two for lower mandible achieved saturation, one for base of mandible brightness, two for cere hue VIS, one for cere brightness and three for rosette hue VIS, and three for rosette hue UV), but the results were qualitatively the same with and without outliers so only the models containing all data are reported.

## Discussion

We replicated Doutrelant et. al.’s (2013) finding that puffins are sexually monochromatic from an avian visual perspective using a much larger sample size (n = 36 vs. n = 162). This supports our underlying assumption that females and males are ornamented to similar degrees.

Our investigation of the characteristics of the puffin’s colorful bill, cere, and rosette yielded conflicting results. The cross-sectional dataset revealed population-level changes in the coloration of multiple regions across the breeding season. In line with this finding, the longitudinal dataset showed that color and brightness perceptibly fluctuate within individuals over the course of the breeding season and are especially dynamic in fleshy structures like the cere and rosette. These analyses support the hypothesis that bill color could act as a signal of quality, with dynamic fluctuations in color potentially corresponding to changes in relative quality during the breeding season. However, we found no relationship between any of our color metrics and a commonly employed proxy of quality, current body condition. A lack of condition-dependence suggests instead that the colorful bill more closely aligns with a signal of identity.

These mixed results led us to further explore the stability of bill color by investigating the time scale over which perceptible color changes occur using our cross-sectional data. We found that the rosette was the only region that consistently became more chromatically perceptible with time, although discriminable changes generally required a longer time interval than indiscriminable changes for all regions. Perhaps unsurprisingly, this fleshy innervated tissue changed on a very rapid time scale (Lucas and Stettenheim 1972; Rosenthal et al., 2012); on average a color change over five days would be sufficient for conspecifics to notice a difference in rosette coloration based on an approximation of puffin visual perception. It is unclear what is driving these color changes in the rosette, as none of the colorimetric variables we investigated (hue VIS, hue UV, achieved saturation) varied according to the time interval between sampling. However, two colorimetric variables (hue VIS and achieved saturation) of the other fleshy structure we examined (cere) did exhibit some evidence of change with sampling interval, such that it became more yellow-orange and less saturated over time. Dynamic changes in the cere and rosette support prior work demonstrating rapid color change in fleshy tissues, such as the 48-hour change in foot color observed in food-deprived blue-footed boobies (*Sula nebouxii*, Velando et al. 2006). The relationship between time and color was nonsignificant for the bill, so we could not calculate a threshold of discriminability, but perceptible changes in bill coloration still occurred within our 21-day time frame. Previous studies have shown that most changes in structures like the keratinized dermal plates of avian bills occur over the course of several weeks (e.g. zebra finches *Taeniopygia guttata,* Blount et al. 2003; European blackbirds *Turdus merula*, Baeta et al. 2008; spotless starlings *Sturnus unicolor*, Navarro et al. 2010), which may partially explain why we were unable to detect a clear relationship in our sample. Yet, avian bills can change rapidly in some species (three days in zebra finches, Ardia et al., 2010; one to three days in goldfinches, Rosenthal et al. 2012), and in our sample some instances of rapid color change occurred (i.e., 4 individuals ≤ 6 days in the upper mandible, 2 individuals ≤ 6 days in the lower mandible). Alternatively, changes in bill or rosette color may be responses to key events such as egg laying or peak food availability, resulting in a more sudden shift that does not vary linearly with time. Regardless of the trajectory of these color changes, any feature capable of fluctuating on the order of days to weeks is too unstable to function as a signal of identity on its own.

The cross-sectional dataset permitted the evaluation of broad trends in color over time across the population, which should not exist if bill color functions as an identity signal. We found several significant changes in colorimetric variables over time; across all regions, hue tended to become more orange-yellow and less UV reflective, while achieved saturation increased and brightness decreased. These results can be juxtaposed with a previous study by Kelly (2015) that examined color change in Atlantic puffin bare part coloration by comparing individuals sampled during the incubation phase (n = 17) and chick rearing phase (n = 17). Kelly (2015) found that saturation in the UV spectrum and brightness of the red and black regions of the mandible were significantly lower in individuals sampled during the chick rearing phase compared to the incubation phase, with no observed differences in hue or saturation in the visual spectrum. While Kelly (2015) investigated a slightly different portion of the breeding season, the observed trends in brightness and UV wavelength contribution are similar between the two studies. Kelly (2015) posits that the observed difference may be partially due to the bill’s function in signaling, such that the bill is more relevant to pair communication earlier in the breeding season. However, we believe this interpretation not only extends beyond the limits of Kelly’s methodology (i.e., cross-sectional), but also fails to recognize more parsimonious explanations.

These cross-sectional trends may instead point to coloration shifts as the end of the breeding season nears and adults prepare to shed their keratinized bill sheaths and reduce the prominence of their rosettes (Harris and Wanless 2011). While an increase in saturation doesn’t typically fit into this narrative, a concurrent shift in hue from red-orange to orange-yellow may explain why achieved saturation increases. Achieved saturation is dependent on the maximum possible saturation value (*r_max_*), which is highest at the photoreceptor vertices (i.e., pure red, pure green, etc) and lowest at the midpoint between the vertices in a tetrahedral color space. Therefore, as the hue shifts from red to orange-yellow, the *r_max_* value decreases and a vector of the same length (i.e., equivalent absolute saturation) would have a higher achieved saturation. Another possibility is that sampling bias occurred, such that only certain individuals remained on the colony and visited their burrows late in the breeding season. These individuals may be low quality if early breeding is advantageous (as in common terns *Sterna hirundo,* Arnold et al. 2004; roseate terns *Sterna dougallii,* Burger et al. 1996; chinstrap penguins *Pygoscelis antarctica,* Moreno et al. 1997; king penguins *Aptendodytes patagonicus,* van Heezik et al. 1994); in this case, their chicks would be less developed than those of high quality individuals and would require more frequent provisioning visits, increasing the likelihood of adult capture in the burrow. In contrast, individuals captured later in the season may be high quality if later breeding is advantageous (i.e., potentially higher synchrony with prey availability, as in thick-billed murres *Uria lomvia,* Gaston et al. 2009; Baird’s sandpiper *Calidris bairdii,* McKinnon et al. 2012; rhinoceros auklets, *Cerorhinca monocerata,* Watanuki et al. 2009), or they are the only individuals with surviving chicks, such that lower quality individuals have already left the colony. Typically, carotenoid-pigmented colors that are more red-shifted, more saturated, brighter, and have less UV reflection are considered “high quality” (e.g., red grouse *Lagopus lagopus scoticus*; Mougeot et al. 2007) and preferred by potential mates or partners in a breeding attempt (e.g., red-legged partridge *Alectoris rufa*, Alonso-Alvarez et al. 2012; reviewed in Hill & McGraw 2006). This would support the first hypothesis that low quality individuals are more likely to be captured later in the breeding season. However, because we did not find a relationship between any metric of color and condition, we cannot draw any definitive conclusions on what aspect of quality may be reflected in the puffin’s bill coloration.

Our study adds to the mixed evidence for condition-dependence of puffin bill and rosette color. Doutrelant (2013) showed that bill coloration was redder (higher value of hue and chroma, lower brightness) for individuals in better body condition and the rosette was more orange (higher values of hue and chroma) for females in better condition. In contrast, Kelly (2015) found no relationship between bill coloration and condition or health. We sought to add clarity to this dispute by employing a dataset containing hundreds of individuals over multiple years. We also used a more repeatable and relevant methodology by 1) directly relating colorimetric variables to body condition, rather than principle components derived from colorimetric descriptors of spectra, and 2) calculating body condition from the residuals of a stepwise linear regression including multiple relevant predictors of mass (Julian date of capture, year, and sex). We ultimately found that none of the examined colorimetric variables were significantly associated with body condition, supporting Kelly’s (2015) conclusion that bill coloration is not condition-dependent. However, this does not exclude the possibility that coloration is related to some other unmeasured aspect of quality uncoupled from our measure of body condition, such as foraging success of high-carotenoid food items, immunocompetence, or oxidative stress. Additionally, while body condition indices are generally good estimates of body fat mass, the index we employed has not been empirically validated, so it is possible that a different measure of body condition is related to coloration (Labocha and Hayes 2012). Another possibility is that bill color is a stronger indicator of quality when selection pressures are high (i.e., years with poor breeding conditions), as seems to be the case in least auklets (*Aethia pusilla*; Jones & Montgomerie, 1992). Nevertheless, the most parsimonious explanation is that bill coloration is not condition-dependent in puffins and that variability in colour is not a meaningful quality signal. The trade-off hypothesis (i.e., one of the primary mechanisms linking carotenoid pigmented features to immune function and health) rests on the assumption that carotenoids are scarce, but recent work shows that carotenoids are generally not a limiting resource physiologically (Simons et al. 2014) There is also an especially weak link between dietary access to carotenoids and bare part coloration in birds (Olson and Owens 2005), so it is unlikely that condition as it relates to foraging ability would be correlated with bill color. Additionally, the mechanisms of carotenoid metabolism vary substantially between and within species, making it difficult to detect relationships within a population, let alone within a species (Svensson and Wong 2011).

Taken together, our results discount the hypothesis that bill coloration signals identity in puffins and provide limited support for bill coloration as a putative signal of quality. The presence of discriminable differences in color over the course of three weeks demonstrates that bill color fluctuates in a perceptible way, but due to a lack of condition-dependence, it remains unknown what these changes represent. Fleshy structures such as the cere and rosette may play an especially important role in signaling because of their ability to change color on a rapid time-scale. Future work should focus on whether color change in the cere and rosette correspond to alternative aspects of quality, such as the ability to provide parental care.

