## Supplemental Tables and Figures for "Bill color is dynamic across the breeding season but not condition-dependent in Atlantic puffins"

Supplementary Tables and Figures

Table S1 Distance PERMANOVA table for chromatic distances between males and females

| Region | Source | Df | SS | MS | F | R^2^ | P |
| --- | --- | --- | --- | --- | --- | --- | --- |
| Upper mandible | Sex | 1 | 14.330 | 14.330 | 3.892 | 0.023 | 0.162 |
|  | Residuals | 167 | 614.941 | 3.682 |  | 0.977 |  |
|  | Total | 168 | 629.271 |  |  | 1.000 |  |
| Lower mandible | Sex | 1 | 1.795 | 1.795 | 0.478 | 0.003 | 0.757 |
|  | Residuals | 167 | 627.476 | 3.757 |  | 0.997 |  |
|  | Total | 168 | 629.271 |  |  | 1.000 |  |
| Cere | Sex | 1 | 0.380 | 0.380 | 0.105 | 0.001 | 0.882 |
|  | Residuals | 167 | 603.363 | 3.613 |  | 0.999 |  |
|  | Total | 168 | 603.743 |  |  | 1.000 |  |
| Rosette | Sex | 1 | 9.319 | 9.319 | 0.366 | 0.002 | 0.757 |
|  | Residuals | 167 | 4248.858 | 25.442 |  | 0.998 |  |
|  | Total | 168 | 4258.177 |  |  | 1.000 |  |

Table S2 Distance PERMANOVA table for achromatic distances between males and females

| Region | Source | Df | SS | MS | F | R^2^ | P |
| --- | --- | --- | --- | --- | --- | --- | --- |
| Upper mandible | Sex | 1 | 5.282 | 5.282 | 2.064 | 0.012 | 0.398 |
|  | Residuals | 167 | 427.307 | 2.559 |  | 0.988 |  |
|  | Total | 168 | 432.589 |  |  | 1.000 |  |
| Lower mandible | Sex | 1 | 0.054 | 0.054 | 0.021 | 0.000 | 0.882 |
|  | Residuals | 167 | 432.535 | 2.590 |  | 1.000 |  |
|  | Total | 168 | 432.589 |  |  | 1.000 |  |
| Base of mandible | Sex | 1 | 22.342 | 22.342 | 5.790 | 0.034 | 0.126 |
|  | Residuals | 167 | 644.358 | 3.858 |  | 0.966 |  |
|  | Total | 168 | 666.700 |  |  | 1.000 |  |
| Cere | Sex | 1 | 3.663 | 3.663 | 1.169 | 0.007 | 0.495 |
|  | Residuals | 167 | 523.446 | 3.134 |  | 0.993 |  |
|  | Total | 168 | 527.109 |  |  | 1.000 |  |
| Rosette | Sex | 1 | 7.029 | 7.029 | 2.002 | 0.012 | 0.398 |
|  | Residuals | 167 | 586.397 | 3.511 |  | 0.988 |  |
|  | Total | 168 | 593.426 |  |  | 1.000 |  |

Table S3 Proportion of discriminable changes in coloration

| ROI | Discriminable | Non-discriminable | Percent Discriminable (%) |
| --- | --- | --- | --- |
| Upper mandible | 25 | 16 | 61.0 |
| Lower mandible | 14 | 27 | 34.1 |
| Cere | 21 | 20 | 51.2 |
| Rosette | 33 | 8 | 80.5 |
| Total (JND = 1) | 93 | 71 | 56.7 |
| Total (JND = 2) | 36 | 128 | 22.0 |
| Total (JND = 3) | 14 | 150 | 8.5 |

Discriminability percentages for distinct regions were calculated at a threshold of JND = 1.

Table S4 Proportion of discriminable changes in brightness

| ROI | Discriminable | Non-discriminable | Percent Discriminable |
| --- | --- | --- | --- |
| Upper mandible | 32 | 9 | 78.0 |
| Lower mandible | 24 | 17 | 58.5 |
| Base of mandible | 30 | 11 | 73.2 |
| Cere | 25 | 16 | 61.0 |
| Rosette | 36 | 5 | 87.8 |
| Total (JND = 1) | 147 | 58 | 71.7 |
| Total (JND = 2) | 87 | 118 | 42.4 |
| Total (JND = 3) | 52 | 153 | 25.4 |

Discriminability proportions for distinct regions were calculated at a threshold of JND = 1.

Table S5 Variation in colorimetric variables with time between sampling dates

| ROI | Response: colorimetric variable | Predictors | Estimate | Std. Error | t | P |
| --- | --- | --- | --- | --- | --- | --- |
| Upper mandible | Hue vis | Sampling interval | 1.72E-04 | 1.25E-03 | 0.137 | 0.892 |
|  |  | Julian date | -2.80E-03 | 1.01E-03 | -2.776 | 0.074(*) |
|  | Hue UV (3 outliers) | Sampling interval | -1.35E-03 | 2.25E-03 | -0.598 | 0.775 |
|  |  | Julian date | 6.78E-03 | 1.73E-03 | 3.926 | 0.014* |
|  | Achieved saturation (2 outliers) | Sampling interval | 4.07E-04 | 2.31E-03 | 0.176 | 0.887 |
|  |  | Julian date | -1.27E-03 | 2.81E-03 | -0.452 | 0.814 |
|  | Brightness (2 outliers) | Sampling interval | -1.02E-02 | 4.69E-03 | -2.167 | 0.126(*) |
|  |  | Julian date | 3.40E-03 | 3.11E-03 | 1.091 | 0.521 |
| Lower mandible | Hue vis | Sampling interval | -4.42E-04 | 1.92E-03 | -0.229 | 0.869 |
|  |  | Julian date | -6.25E-04 | 1.55E-03 | -0.403 | 0.814 |
|  | Hue UV (1 outlier) | Sampling interval | -2.89E-03 | 3.00E-03 | -0.963 | 0.544 |
|  |  | Julian date | 7.66E-03 | 3.06E-03 | 2.503 | 0.107(*) |
|  | Achieved saturation (3 outliers) | Sampling interval | -3.90E-03 | 1.87E-03 | -2.089 | 0.126(*) |
|  |  | Julian date | -1.09E-03 | 1.79E-03 | -0.608 | 0.775 |
|  | Brightness | Sampling interval | -1.07E-03 | 2.99E-03 | -0.358 | 0.816 |
|  |  | Julian date | 2.38E-03 | 2.41E-03 | 0.985 | 0.544 |
| Mandible base | Brightness (2 outliers) | Sampling interval | -5.70E-03 | 2.37E-03 | -2.402 | 0.108(*) |
|  |  | Julian date | 7.28E-04 | 1.58E-03 | 0.461 | 0.814 |
| Cere | Hue vis (3 outliers) | Sampling interval | 0.28 | 0.13 | 2.153 | 0.126(*) |
|  |  | Julian date | 0.013 | 6.79E-03 | 1.852 | 0.182 |
|  |  | Sampling interval x Julian date | -1.42E03 | 6.67E-04 | -2.147 | 0.126(*) |
|  | Hue UV | Sampling interval | -6.60E-04 | 2.42E-03 | -0.273 | 0.860 |
|  |  | Julian date | 1.78E-03 | 1.95E-03 | 0.913 | 0.559 |
|  | Achieved saturation (3 outliers) | Sampling interval | -6.10E-03 | 2.16E-03 | -2.819 | 0.074(*) |
|  |  | Julian date | 5.22E-03 | 1.86E-03 | 2.805 | 0.074(*) |
|  | Brightness (2 outliers) | Sampling interval | -8.04E-03 | 3.82E-03 | -2.106 | 0.126(*) |
|  |  | Julian date | -3.53E-03 | 2.54E-03 | -1.392 | 0.379 |
| Rosette | Hue vis | Sampling interval | -2.54E-03 | 1.83E-03 | -1.388 | 0.379 |
|  |  | Julian date | -1.84E-03 | 1.47E-03 | -1.249 | 0.452 |
|  | Hue UV | Sampling interval | 2.78E-03 | 1.35E-03 | 2.054 | 0.126(*) |
|  |  | Julian date | 1.10E-03 | 1.09E-03 | 1.011 | 0.544 |
|  | Achieved saturation (3 outliers) | Sampling interval | -3.45E-03 | 2.90E-03 | -1.192 | 0.469 |
|  |  | Julian date | 1.32E-03 | 3.07E-03 | -0.430 | 0.814 |
|  | Brightness | Sampling interval | -2.48E-02 | 1.00E-02 | -2.465 | 0.107(*) |
|  |  | Julian date | -3.17E-03 | 8.10E-03 | -0.392 | 0.814 |

­*P* values were corrected with the false discovery rate method. (*) indicates significance prior to the false discovery rate correction, and * indicates that the variable remained significant after applying the correction.

Table S6 Relationship between colorimetric variables and body condition

| ROI | Response variable | Estimate | Std. Error | t | P |
| --- | --- | --- | --- | --- | --- |
| Upper mandible | Hue VIS | 2.87E-04 | 1.77E-04 | 1.620 | 0.85 |
|  | Hue UV | 1.10E-04 | 3.33E-04 | 0.329 | 0.97 |
|  | Achieved saturation | 9.01E-06 | 2.86E-04 | 0.031 | 0.99 |
|  | Brightness | 4.99E-06 | 3.13E-04 | 0.016 | 0.99 |
| Lower mandible | Hue VIS | 4.63E-04 | 1.98E-04 | 2.344 | 0.35(*) |
|  | Hue UV | -2.49E-04 | 3.24E-04 | -0.769 | 0.94 |
|  | Achieved saturation | 2.45E-04 | 2.95E-04 | 0.829 | 0.94 |
|  | Brightness | -1.82E-05 | 3.17E-04 | -0.057 | 0.99 |
| Base of mandible mandible | Brightness | 2.11E-04 | 1.80E-04 | 1.171 | 0.85 |
| Cere | Hue VIS | -1.03E-04 | 1.95E-04 | -0.530 | 0.96 |
|  | Hue UV | 2.23E-04 | 2.87E-04 | 0.779 | 0.94 |
|  | Achieved saturation | 5.12E-04 | 4.25E-04 | 1.205 | 0.85 |
|  | Brightness | 1.38E-04 | 3.78E-04 | 0.365 | 0.97 |
| Rosette | Hue VIS | -1.10E-04 | 2.20E-04 | -0.498 | 0.96 |
|  | Hue UV | 1.77E-04 | 1.53E-04 | 1.156 | 0.85 |
|  | Achieved saturation | 2.07E-04 | 3.57E-04 | 0.580 | 0.96 |
|  | Brightness | 4.99E-06 | 3.13E-04 | 0.016 | 0.99 |

Statistical significance at α = 0.05. (*) indicates a significant *P* value before application of the false discovery rate.

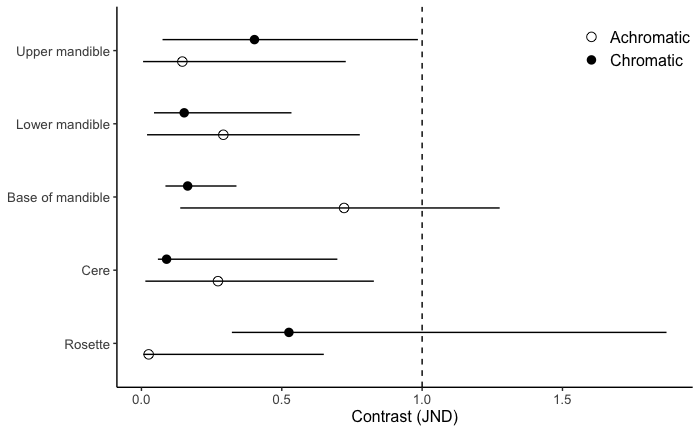

Figure S1

Bootstrapped 95% C.I.’s for mean chromatic and achromatic distances between males and females in color space, grouped by region of the bill

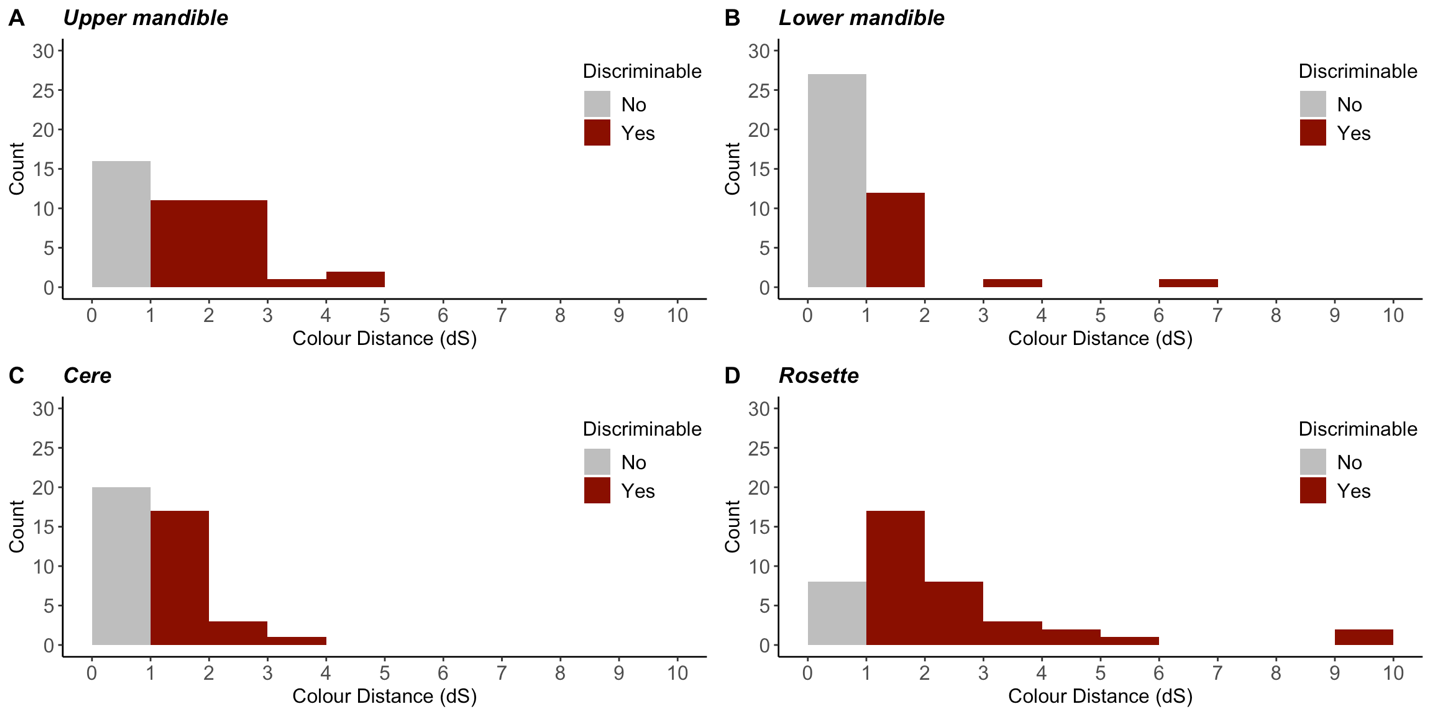

Figure S2

Distribution of color distances between patches sampled within individuals for the A) upper mandible, B) lower mandible, C) cere, and D) rosette.

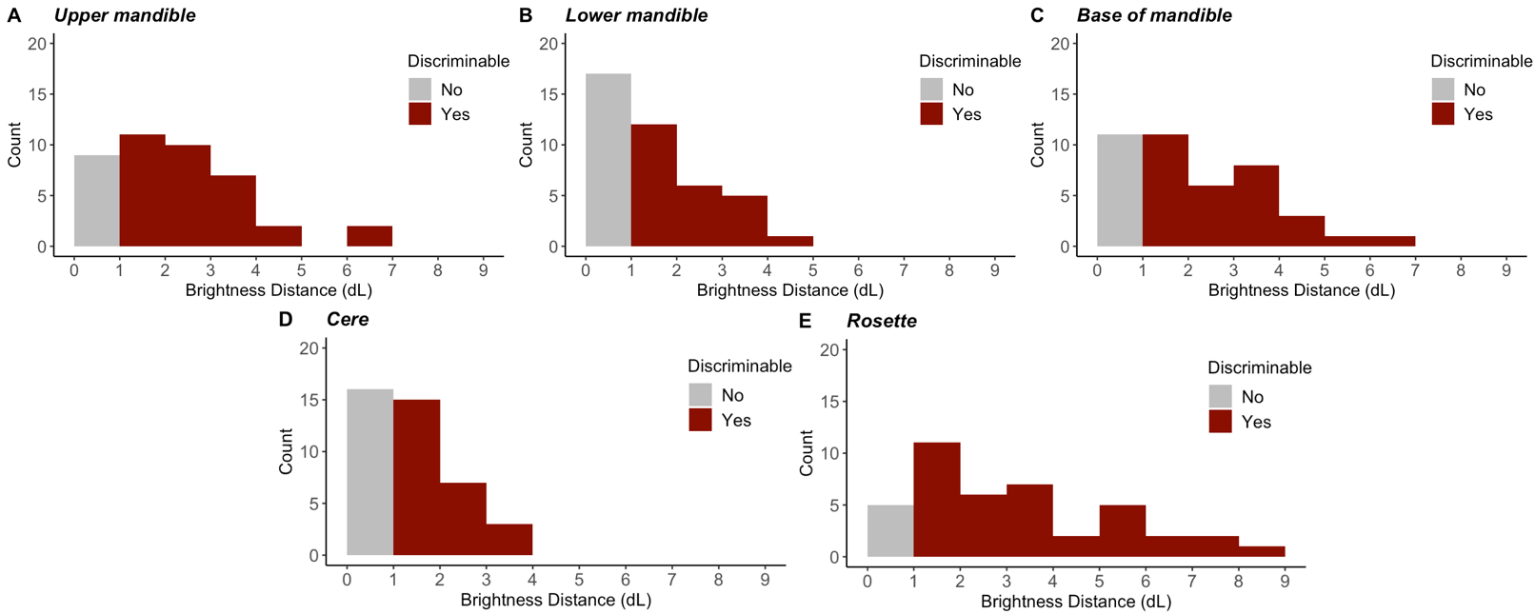

Figure S3

Distribution of brightness distances between patches sampled within individuals for all five regions of interest: A) upper mandible, B) lower mandible, C) base of mandible, D) cere, and E) rosette
